# A systematic evaluation of *Mycobacterium tuberculosis* Genome-Scale Metabolic Networks

**DOI:** 10.1101/837401

**Authors:** Víctor A López-Agudelo, Emma Laing, Tom A Mendum, Andres Baena, Luis F Barrera, Dany JV Beste, Rigoberto Rios-Estepa

## Abstract

The metabolism of the causative agent of TB, *Mycobacterium tuberculosis* (Mtb) has recently re-emerged as an attractive drug target. A powerful approach to study Mtb metabolism is to use a systems biology framework, such as a Genome-Scale Metabolic Network (GSMN) that allows the dynamic interactions of the many individual components of metabolism to be interrogated together. Several GSMNs networks have been constructed for Mtb and used to study the complex relationship between Mtb genotype and phenotype. However, their utility is hampered by the existence of multiple models of varying properties and performances. Here we systematically evaluate eight recently published metabolic models of Mtb-H37Rv to facilitate model choice. The best performing models, sMtb2018 and iEK1011, were refined and improved for use in future studies by the TB research community.

## Introduction

*Mycobacterium tuberculosis* (Mtb) is the causative bacterium agent of the global tuberculosis (TB) epidemic, which is now the biggest infectious disease killer worldwide, causing 1.6 million deaths in 2017 alone [1]. Mtb is an unusual bacterial pathogen, as it is able to cause both acute life threatening disease and a clinically latent infections that can persist for the lifetime of the human host [2, 3]. Metabolic reprogramming in response to the host niche during both the acute and the chronic phase of TB infections is a crucial determinant of virulence [4–7]. With the worldwide spread of multi- and extensively-resistant strains of Mtb thwarting the control of this global emergency, new drugs against Mtb are urgently needed and metabolism presents an attractive target [8, 9].

Genome-scale constraint-based modelling has proved to be a powerful method to probe the metabolism of Mtb. The first Genome Scale Metabolic Networks (GSMNs) of Mtb were published in 2007 by Beste (GSMN-TB) [10] and Jamshidi (iNJ661) [11] and have been used as a platform for interrogating high throughput ‘omics’ data, by simulating bacterial growth, generating hypothesis and drug discovery and are also the backbone for subsequent model iterations [12–20]. In 2014, Rienksma and colleagues developed a genome-scale model of Mtb metabolism (sMtb) building upon three previously published models [15]. More recently Rienksma and colleagues utilized RNA sequence data to provide condition-specific biomass reactions and identify metabolic drug responses during Mtb infection of human macrophages [21, 22] thus providing a model more suited to exploring the metabolism of Mtb during human infection. In 2018 the first consolidated genome-scale metabolic network, iEK1011 was constructed using standardized nomenclature of metabolites and reactions from the BiGG model database [23, 24]. The utility of Mtb GSMNs will continue to increase, as more biological data is published and existing models are updated [25].

With so many well-annotated GSMN’s of Mtb available (Fig 1), a crucial first step in any genome scale exploration of the metabolism of Mtb is the selection of an appropriate model. Here, we systematically evaluate the performance of eight recently published GSMN of Mtb-H37Rv. In addition to comparing the metrics of the models descriptively in terms of size, connectivity, number of blocked reactions and gaps in the network, we also identify the thermodynamically infeasible, and energy generating cycles that could significantly impact on the accuracy of flux simulations. Using Flux Balance Analysis (FBA) we performed growth analysis and compared the models’ ability to predict gene essentiality when grown on different carbon and nitrogen sources including growth on cholesterol, which is thought to be the major carbon source for intracellular Mtb within its human host macrophage.

**Fig 1.**
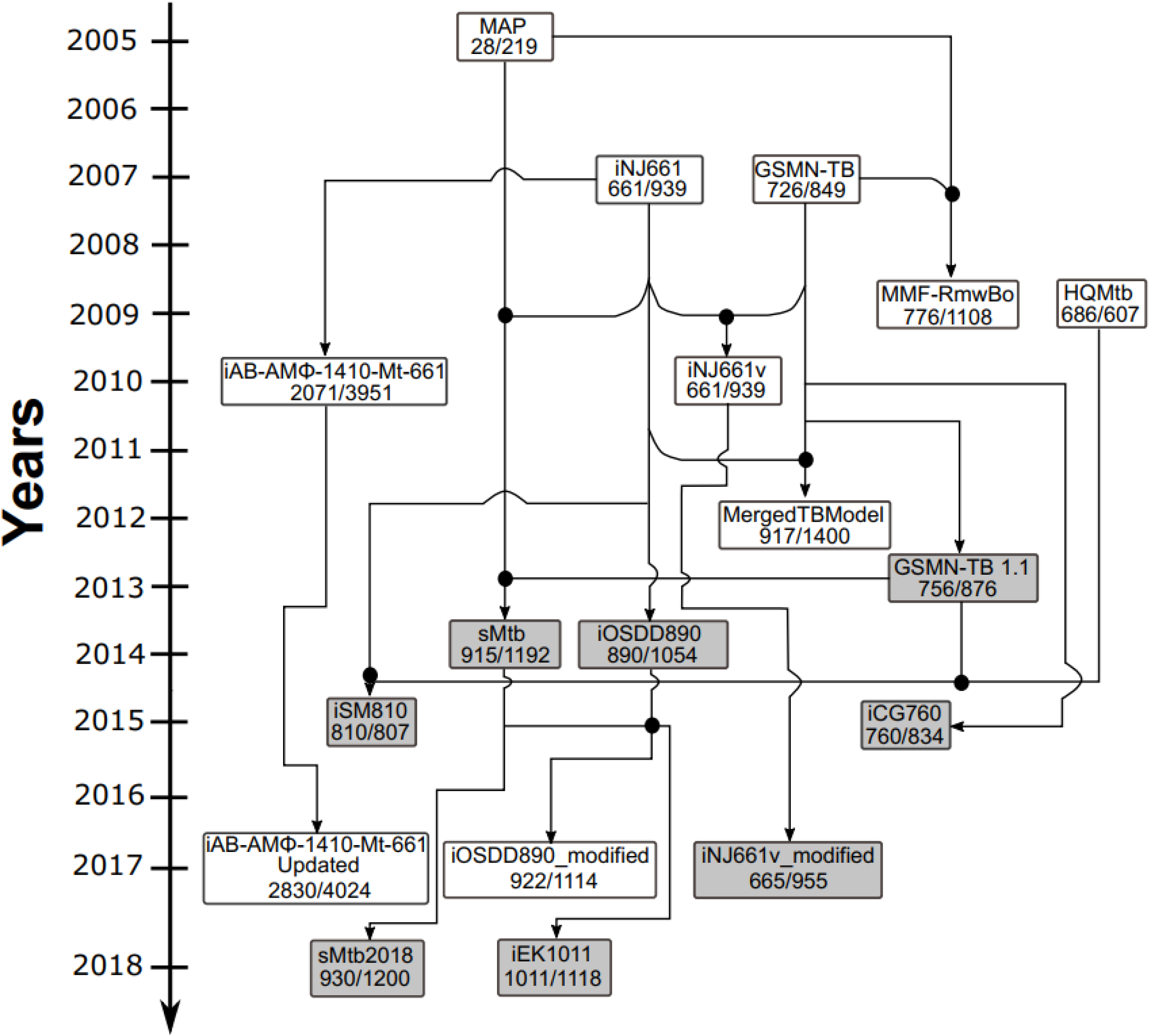
The evolution of Genome Scale Metabolic models of Mtb. Models highlighted in gray were analysed in this study. Numbers denote genes/intracellular reactions. Black circles are indicative of merged Mtb models.

This work provides an inventory of the available GSMN-TB and their utility in recapitulating aspects of Mtb metabolism. In addition, we present updated versions of the highest performing models iEK1011 and sMtb2018 (iEK1011_2.0 and sMtb2.0) which provide multi-scale simulation platforms available for the TB research community and will therefore inform future studies on the metabolism of this deadly pathogen.

## Results and Discussion

### Descriptive evaluation of the models

Each of the GSMNs analysed in this study (Fig 1, S1 Appendix) combine knowledge from genome annotations, literature and measured biochemical compositions of Mtb. The complex linkage between genotype and phenotype is made by gene-protein-reaction (GPR) associations, implemented as Boolean rules in order to connect gene functions to enzyme complexes, isozymes or promiscuous enzymes, and then to biochemical reactions [26]. Using set theory, we computed the intersection between all sets of the model’s genes (Fig 2, and S1 Table). In accordance with expectations, the pairwise matrix (Fig 2) demonstrates that Mtb models constructed from the same ancestor (iNJ661 or GSMN-TB), are more similar (Fig. 1, Fig. 2). By contrast the consolidated models iEK1011 and sMtb2018 share gene similarities (>60%, <85% for iEK1011; and >60%, <98.4%) with all the other models demonstrating an independence from iNJ661 and GSMN-TB.

**Fig 2.**
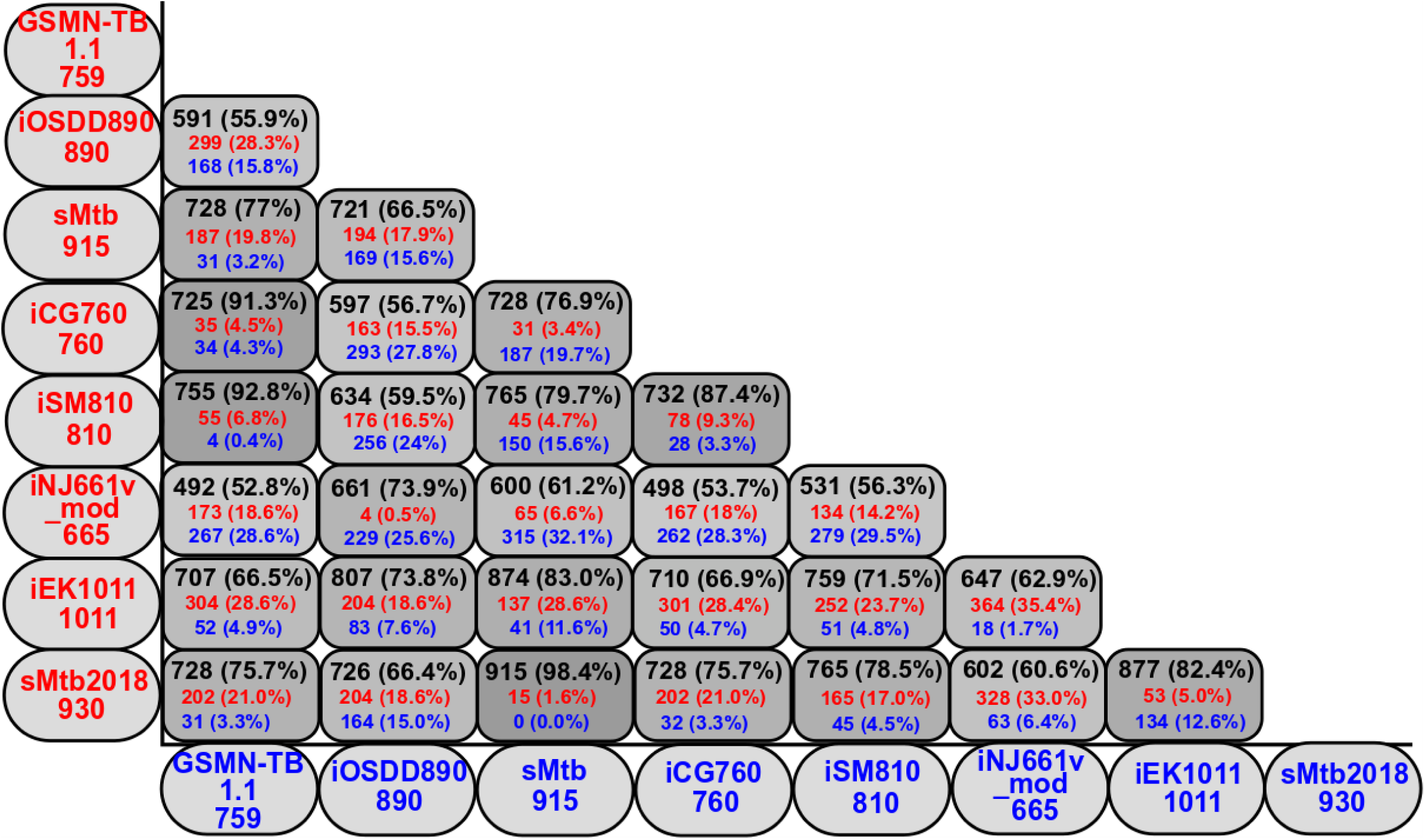
Pairwise matrix of shared genes among Mtb models. Values in black (genes in common between the models), blue (genes specific to the model indicated), and red (genes specific to the model indicated) represent the number and percentage of specific model genes specified in the y- and x-axis.

All the models contain essential metabolic pathways such as carbon, nitrogen, nucleotides, and cofactor metabolism (S2 Table), encoded by 479 common genes that can be used to construct a core metabolic network for Mtb [27]. The models sMtb, sMtb2018 and iEK1011 had the greatest coverage of GPR associations and contain other genes associated with survival and virulence within the host such as transport, respiratory chain, fatty acid, dimycocerosate esters and mycobactin metabolism (S3 Table) and are therefore good candidates to study Mtb metabolism during intracellular growth [28, 29].

In contrast, iOSDD890 (Fig 1) contains a high percentage of genes associated with nitrogen, propionate, pyrimidine, peptidoglycan, pyruvate and cofactor metabolism, but has a lower percentage of genes associated with glycerophospholipid metabolism, cholesterol degradation and fatty acid biosynthesis. Likewise, the iNJ661v_modified model (Fig 1) has poor annotation of genes involved in lipid-metabolism (e.g. β-oxidation, cholesterol degradation, fatty acid biosynthesis, lipid biosynthesis and mycolic acid biosynthesis). These models therefore have limitations for *in silico* simulation of Mtb growing on these physiologically relevant lipid sources and for modelling growth *in vivo*.

### Checking mass and charge balances of biochemical reactions

Currency metabolites like water, protons, ATP, and cofactors like NADH, NADPH, FADH2, CoA, etc. are ubiquitous and essential for metabolism. The addition of these cofactor metabolites in GSMNs and in the biomass reaction considerably improves phenotype predictions and is a hallmark of good quality reconstructions [19, 30]. In order to check this we converted the GSMNs into substance graphs (using a local script) where metabolites (nodes) are connected by edges (undirected and unweighted) if they appear in the same reaction [31] and computed node degrees (number of edges connected to the node) (Table 2, and S4 Table),

**Table 2.**
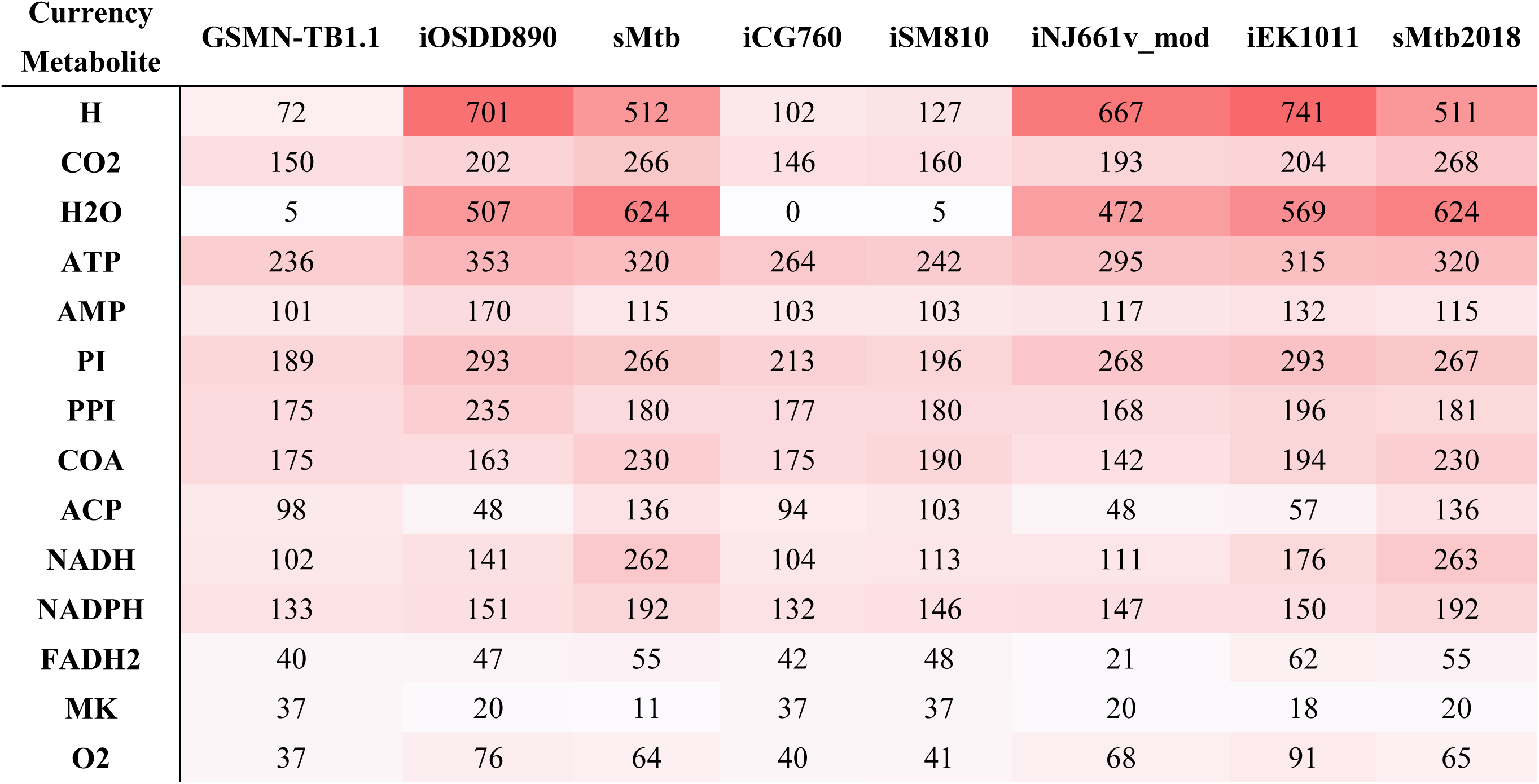
Degree values for currency metabolites of Mtb GSMNs. PI: Phosphate, PPI: diphosphate, ACP: acyl-carrier protein, MK: menaquinone.

| Currency Metabolite | GSMN-TB1.1 | iOSDD890 | sMtb | iCG760 | iSM810 | iNJ661v_mod | iEK1011 | sMtb2018 |
| --- | --- | --- | --- | --- | --- | --- | --- | --- |
| **H** | 72 | 701 | 512 | 102 | 127 | 667 | 741 | 511 |
| **CO2** | 150 | 202 | 266 | 146 | 160 | 193 | 204 | 268 |
| **H2O** | 5 | 507 | 624 | 0 | 5 | 472 | 569 | 624 |
| **ATP** | 236 | 353 | 320 | 264 | 242 | 295 | 315 | 320 |
| **AMP** | 101 | 170 | 115 | 103 | 103 | 117 | 132 | 115 |
| **PI** | 189 | 293 | 266 | 213 | 196 | 268 | 293 | 267 |
| **PPI** | 175 | 235 | 180 | 177 | 180 | 168 | 196 | 181 |
| **COA** | 175 | 163 | 230 | 175 | 190 | 142 | 194 | 230 |
| **ACP** | 98 | 48 | 136 | 94 | 103 | 48 | 57 | 136 |
| **NADH** | 102 | 141 | 262 | 104 | 113 | 111 | 176 | 263 |
| **NADPH** | 133 | 151 | 192 | 132 | 146 | 147 | 150 | 192 |
| **FADH2** | 40 | 47 | 55 | 42 | 48 | 21 | 62 | 55 |
| **MK** | 37 | 20 | 11 | 37 | 37 | 20 | 18 | 20 |
| **O2** | 37 | 76 | 64 | 40 | 41 | 68 | 91 | 65 |

From this analysis the degree values indicated that GSMN-TB 1.1, iCG760 and iSM810 models have the lowest participation of currency metabolites (Table 2). Water and protons were the most underrepresented metabolites (low degree values), indicating that these models may not be correctly balanced. It is important that biochemical reactions are charge and mass balanced. Unbalanced reactions in GSMNs may allow proton or ATP production out of nothing [32]. In order to test whether the Mtb GSMNs are mass and charge balanced, we used the COBRA Toolbox function “checkMassChargeBalance” [33]. Unfortunately, we were not able to perform this analysis for GSMN-TB1.1, iCG760 and iSM810 due to the lack of standard metabolite formulas in these models (Table 3). iEK1011 has the lowest number of imbalanced reactions (4) compared with sMtb2018 (8), sMtb, (12), iNJ661v_modified (13) and iOSDD890 (78). The majority of the imbalanced reactions belonged to cell wall biosynthetic pathways, including arabinogalactan, peptidoglycan, and mycolic acid biosynthesis (S5 Table) reflecting the difficulties in rebuilding accurate metabolite formula for complex cell wall components.

**Table 3.** Global features of the Mtb metabolic models analyzed in this study. DEM: Dead-End Metabolite, Metabolites wo F: Metabolites without formula, Imbal. Reactions: Imbalanced Reactions, URs: Unbounded Reactions, TICs: Thermodynamically Infeasible Cycles, diss. Flux: dissipation flux, MW: Molecular Weight, UD: Undetermined.

|  | GSMN-TB 1.1 | iOSDD890 | sMtb | iCG760 | iSM810 | iNJ661v_mod | iEK1011 | sMtb2018 |
| --- | --- | --- | --- | --- | --- | --- | --- | --- |
| **Reactions** | 876 | 1152 | 1311 | 965 | 938 | 1054 | 1228 | 1321 |
| **Intracellular Reactions** | 876 | 1055 | 1192 | 864 | 938 | 956 | 1118 | 1200 |
| **Metabolites** | 667 | 961 | 1047 | 754 | 724 | 840 | 998 | 1049 |
| **DEMs** | 25 | 162 | 33 | 84 | 53 | 90 | 110 | 34 |
| **Blocked Reactions** | 98 (11%) | 290 (25%) | 92 (7%) | 89 (9%) | 117 (12%) | 153 (14%) | 138 (11%) | 92 (7%) |
| **Metabolites wo F** | 667 | 0 | 2 | 754 | 724 | 0 | 4 | 2 |
| **Imbal. Reactions** | UD | 78 | 12 | UD | UD | 13 | 4 | 8 |
| **URs** | 27 (3%) | 52 (5%) | 70 (6%) | 45 (5%) | 33 (3.5%) | 67 (7%) | 20 (2%) | 75 (6%) |
| **TICs** | 8 | 17 | 16 | 8 | 8 | 23 | 4 | 17 |
| **ATP diss. Flux** | 0 | 0 | 0 | 0.33 | 0 | 0 | 0 | 0 |
| **GTP diss. Flux** | 0 | 0 | 0 | 0.33 | 0 | 0 | 0 | 0 |
| **CTP diss. Flux** | 0 | 0 | 0 | 0.33 | 0 | 0 | 0 | 0 |
| **UTP diss. Flux** | 0 | 0 | 0 | 0.33 | 0 | 0 | 0 | 0 |
| **Biomass MW** | UD | 1.0070 | 1.0126 | UD | UD | 1.0094 | 1.0937 | 1.0126 |

Another potential error in GSMNs is the molecular weight of biomass, which should be defined as 1 g/mmol. Discrepancy in biomass weight can arise as a result of unbalanced reactions which impacts on the reliability of flux predictions using Flux Balance Analysis (FBA). This affect can be amplified when host-pathogen interactions are simulated by integrating host and pathogen metabolic models [34]. Chang and colleagues (2017) developed a systematic algorithm for checking if the biomass molecular weight from GSMNs deviates from 1 g/mmol,. Using this algorithm [34] we found deviations of <10% from 1 g/mmol in all the Mtb models tested (Table 3) (iOSDD890 (0.7%), iNJ661v_mod (0.9%), sMtb (1.2%), sMtb2018 (1.2 %) and iEK1011 (9%)) demonstrating that these models are suitable for modelling the metabolism of Mtb within the host.

### Blocked reactions and dead-end metabolites

Identifying blocked reactions within a GSMN is important for identifying metabolic dead zones caused by dead-end metabolites (metabolites that are not consumed) [35–37]. Using the MC3 algorithm [38], we show that Mtb models derived from GSMN-TB (GSMN-TB1.1, iCG760, and iSM810) have a smaller number of blocked reactions in comparison with the iNJ661 derived models (iNJ661v_modified, and iOSDD890) (Table 3). sMtb and sMtb2018 have the lowest percentage (7%) of blocked reactions, in contrast to iOSDD890, which has the highest percentage (25%). All the Mtb genome-scale models included blocked reactions in lipid, cofactor, sugar and amino acid metabolism; iOSDD890, iSM810 and iNJ661v_mod had blockages in important pathways such as glycolysis and redox metabolism (S6 Table). Most of the models excluding iCG760 and iSM810 contained gaps in the vitamin B12 pathway [39]. Specifically, we found that aqua(III) cobalamin and different cobalt-precorrins were not connected by reactions in most of the networks. This cofactor is necessary for activation of essential pathways such as nucleotide, propionate, and amino acids metabolism [39]. However, the existence of a functional B12 biosynthetic pathway is still under debate.A possible transporter of vitamin B12 has been reported [40], but there remains no direct evidence of Mtb scavenging vitamin B12 from its intracellular niche [41, 42].

Of those models that contained a pathway for cholesterol degradation (GSMN-TB1.1, iCG760, iSM810, iEK1011, sMtb, and sMtb2018). The GSMN-TB1.1 cholesterol degrading pathway contained a number of dead end metabolites making this model unsuitable for exploring the metabolism of this important *in vivo* carbon source.

### Thermodynamic and energetic properties

Integrating thermodynamics data into GSMNs is extremely useful in order to check the feasibility of reactions and their directionality [43, 44]. Although, Mtb GSMNs have been built from thermodynamics information, current Mtb GSMNs have never been checked for infeasible internal flux cycles (also named type III pathways), which are reactions that do not exchange metabolites with the surroundings and therefore violate the second law of thermodynamics [43,45,46]. A tractable way to identify reactions participating in these thermodynamically infeasible cycles (TICs) is by defining the set of reactions required for an unbounded metabolic flux under finite or even zero substrate uptake inputs. Using Flux Variability Analysis (FVA) the Unbounded Reactions (URs) can be identified as those reactions with fluxes at the upper and/or lower bound constraints. Thus we identified the thermodynamically infeasible cycles (TIC) using a local script following methodologies based on FVA and the analysis of the null space of the stoichiometric matrices (S7 Table) [44, 47]. Using this approach we show that models descended from GSMN-TB (GSMN-TB1.1, iCG760, and iSM810) have a lower percentage of unbounded reactions as compared with iNJ661 ancestors (iSM810, iNJ661v_modified). Interestedly, the sMtb2018 model has an increased number of unbounded reactions as compared to the original sMtb (Table 3).

Fritzemeier and colleagues demonstrated that over 85% of genome-scale models that lack exhaustive manual curation contain Energy Generating Cycles (EGCs) [48, 49]. These cyclic net fluxes are entirely independent of nutrient uptakes (exchange fluxes) and therefore have a substantial effect on the predictions of constraint-based analyses, as they basically generate energy out of nothing. Using FBA with zero nutrient uptake [49] but maximizing energy dissipation reactions for ATP, GTP, CTP and UTP we show that iCG760 is (Table 3), the only Mtb genome-scale model that contains EGCs.

### Gene essentiality metrics

An effective and commonly employed predictive matrix for GSMNs is the ability to reproduce high throughput gene essentiality data [50]. Several high throughput transposon mutagenesis screens have been performed for Mtb [51–57] in different *in vitro* conditions. To compare our models we used a transposon insertion sequence dataset produced by Griffin et al [54] in which genes that were differentially required for growth on cholesterol as compared with glycerol [54] were identified. Cholesterol is an important intracellular source of carbon when Mtb is growing within its host and cholesterol metabolism has been highlighted as a potential drug target [58]. Whilst Griffin and colleagues identified differentially required genes, this data was not used to identify the genes required for growth on cholesterol as an individual condition. We reanalyzed the Griffin’s transposon sequencing data with the statistical Bayesian/Gumbel Method incorporated into the software TRANSIT [59], to identify genes required for growth on glycerol, or growth on cholesterol (S8 and S9 Tables). Only genes categorized as essential (ES) and non-essential (NE) were considered for this analysis.

We evaluated the overall predictive power of all the Mtb GSMNs versus four high throughput gene essentiality data [54,56,57] by computing the Area Under the Curve (AUC) of the Receiver Operating Characteristic (ROC) (S1 Fig). The predictive power of the six GSMNs which contain the cholesterol degradation pathway (GSMN-TB1.1, iCG760, iSM810,sMtb, sMtb2018, and iEK1011) showed that for both cholesterol and glycerol minimal media the models derived from GSMN-TB [10] as a core metabolic network have better predictive capacities than those using iNJ661 [11] (S10 and S11 Tables). However the recently curated model iEK1011 had the highest predictive capability overall. The supremacy of iEK1011 also was confirmed by comparing the predictive power of the models for Mtb grown in standard Middlebrook 7H9 OADC [56] data (S1C Fig and S12 Table).

We further used the essentiality dataset generated by Minato and colleagues who identified conditionally essential genes on different *in vitro* medium including a rich medium called “MtbYM”, which contains several carbon sources, nitrogen sources, amino acids, nucleotide bases, cofactors, and other nutrients [57]. Although this media supported growth similarly to 7H9 medium, the overall predictive power of the Mtb GSMNs in this medium (S1D Fig amd S13 Table) was 10% lower compared to the other media (S1A-S1C Figs), probably because biomass objective functions of the Mtb GSMNs ancestors were built and validated using growth on 7H9, 7H10, Youman’s, CAMR, and Roisin’s minimal medium [10, 11]. However these analyses were able to demonstrate the power of ability of these models to accurately predict gene essentiality even under un-anticipated nutritional conditions.

We identified the genes that all of the GSMN’s were unable to correctly assign essentiality (S14 Table, Fig 3). Using a fixed threshold value of 5% of the maximum wild-type growth rate (WTGR) we identified *in silico* essential and non-essential genes and using experimental high throughput gene essentiality data identified True Positives (TP), True Negatives (TN), False Positives (FP-a gene which is essential for in silico growth but non-essential by Tn-seq) and False Negatives (FN-in silico the gene is non-essential but the biological data predicts essentiality). FN genes include genes known to have a major role in Mtb central carbon metabolism e.g., *icl* (Rv0467, isocitrate lyase), *glt*A (Rv0896, Citrate synthase), *glp*D2 (Rv3302c, Glycerol-3-phosphate dehydrogenase), *pyk* (Rv1617, Pyruvate Kinase), *suc*C and *suc*D (Rv0951, Rv0952, Succinyl-CoA ligase), among others (S14A Table). Some of these genes e.g. *icl* and *glt*A are considered conditionally essential genes in the Online GEne Essentiality (OGEE) database [60], because they are classified as NE genes in 7H10 medium but ES in minimal medium [53, 55]. This may reflect the presence of alternative routes *in silico* that are not feasible *in vivo* due to regulatory constraints. However, they may also reflect inaccuracies in the predictive abilities of transposon mutagenesis studies. Some of the FP genes are involved in mycolic acid biosynthesis (S14B Table), e.g., *mma*A2 (Rv0644c, Cyclopropane mycolic acid synthase), and *mas* (Rv2940c, mycocerosic acid synthase). These genes were inaccurately classified as ES *in silico* but experimentally as NE. This is a direct reflection of our incomplete knowledge of these mycolate producing pathways and the essentiality of the various mycolates produced.

**Fig 3.**
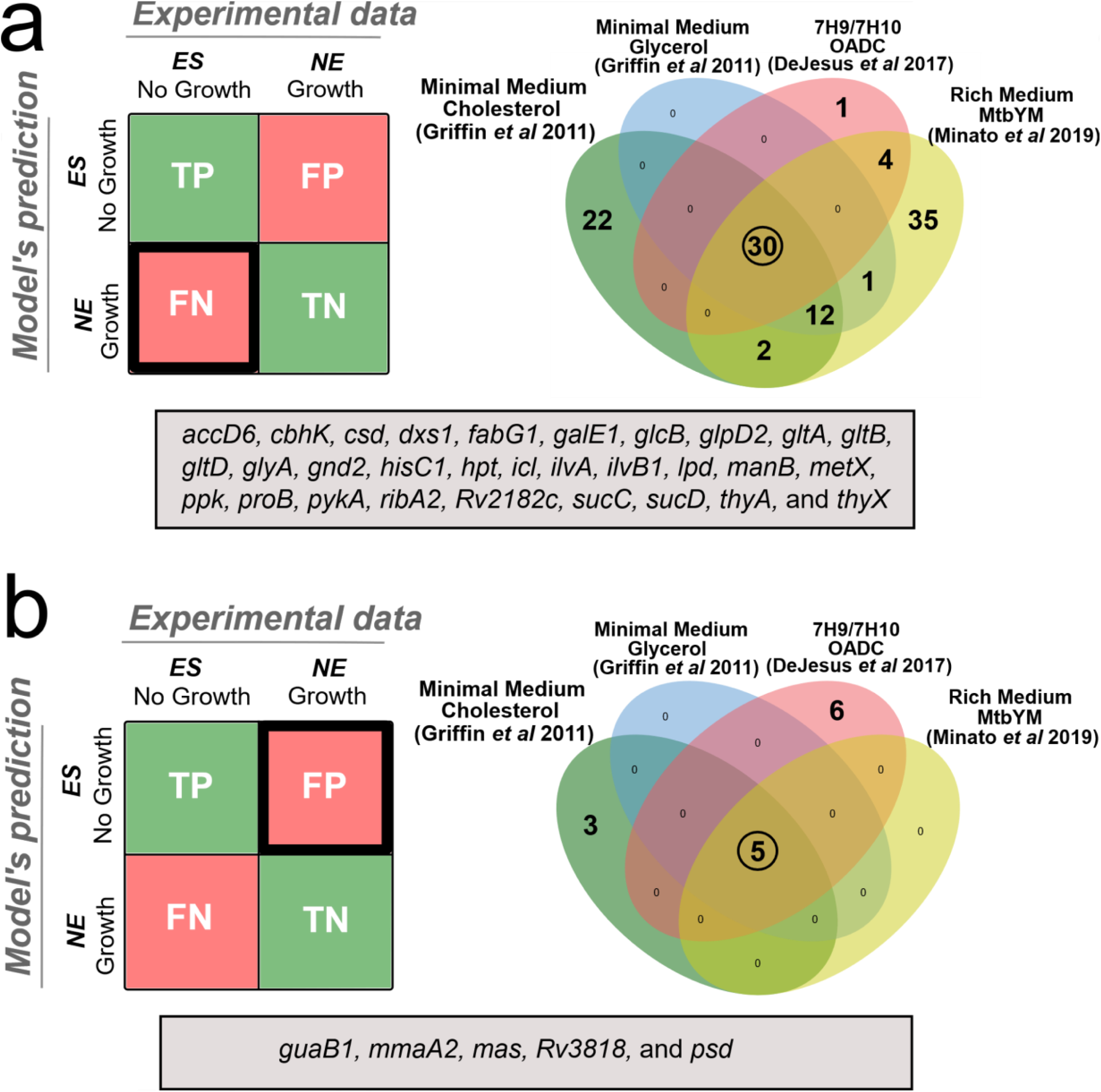
False Negative and False Positive predictions in the evaluated media. (a) Venn diagram for predicted false negative (FN) genes; (b) Venn diagram for predicted false positive (FP) genes. Genes in the gray box represents the intersection of all FN and FP genes in the four media. Genes are classified as True-positives (TP) if model simulation predicts no growth when essential genes are deleted, False-positives (FP) if model simulation predicts no growth when not essential genes are deleted, True negatives (TN) if model simulation predicts growth when not essential genes are deleted and False negatives (FN) if model simulation predicts growth when essential genes are deleted.

### Growth metrics

Mtb is able to metabolise several carbon and nitrogen sources both *in vitro* and when growing in the host [61–64], and therefore we evaluated the growth metrics of Mtb GSMNs on 30 sole carbon and 17 sole nitrogen sources (Fig 4). The *in silico* results were compared with available *in vitro* experimental data from Biolog Phenotype microarray and minimal media data [14, 65]. Interestingly the recent consolidated models, iEK1011 and sMtb, had the poorest performance of all the models in predicting growth of Mtb in unique carbon and nitrogen sources (Fig 4A and 4B). A fundamental issue with the Mtb models descended from iNJ661 is that they all require glycerol for growth as this is present in the biomass formulation. Both iEK1011 and sMtb were unable to grow *in silico* on cholesterol, acetate, oleate, palmitate and propionate on sole carbon sources. We hypothesise that this is as a result of inaccuracies in the reactions associated with redox metabolism and oxidative phosphorylation and specifically including menaquinone-dependent reactions such as fumarate reductase and succinate dehydrogenase would correct this anomalie. To test this hypothesis we added a menaquinone-dependent succinate dehydrogenase reaction into sMtb (Q[c] + SUCC[c] -> QH2[c] + FUM[c]). In support of our hypothesis this corrected the *in silico* growth phenotype of Mtb growing on acetate, cholesterol, propionate and fatty acids (Fig 4A and S15 Table). Although iEK1011 also contains a fumarate reductase reaction that is linked to menaquinone/demethylmenaquinone, it does not contain a menoquinone-dependent succinate dehydrogenase (the reverse reaction). As was the case for sMtb, the addition of a new menaquinone-dependent succinate dehydrogenase reaction (mqn8[c] + succ[c] -> fum[c] + mql8[c]) to iEK1011 significantly improves its growth predictions on sole carbon sources (S15I and S15J Tables). These simulations are supported by experimental data demonstrating that fumarate reductase and succinate dehydrogenase are essential for the survival of Mtb in glycolytic and non-glycolytic substrates [66, 67]. Succinate dehydrogenase is a bifunctional enzyme that is part of the TCA cycle and complex II of the electron transport chain, coupling the oxidation of succinate to fumarate, with the corresponding reduction of membrane-localized quinone electron carriers [67, 68]. Mtb has multiple succinate dehydrogenases and fumarate reductases that are essential for the survival of Mtb survival during hypoxia [66,67,69–71]. Succinate is central to much of Mtb’s lipid metabolism: host derived cholesterol, uneven chain length fatty acids or methyl branched amino acids all generate propionyl-CoA that can be channeled into the methylcitrate cycle to produce succinate (S2 Fig), while acetyl-CoA produced by β-oxidation of host derived even-chain fatty acids is metabolized through the glyoxylate shunt to also produce succinate (S2 Fig). Succinate oxidation by succinate dehydrogenases is therefore a critical step, as the enzyme couples the TCA cycle with electron transport chain and oxidative phosphorylation [69]. Having multiple succinate dehydrogenases provides Mtb with the metabolic flexibility to survive within the human host where the REDOX and nutritional environment are liable to vary.

**Figure 4.**
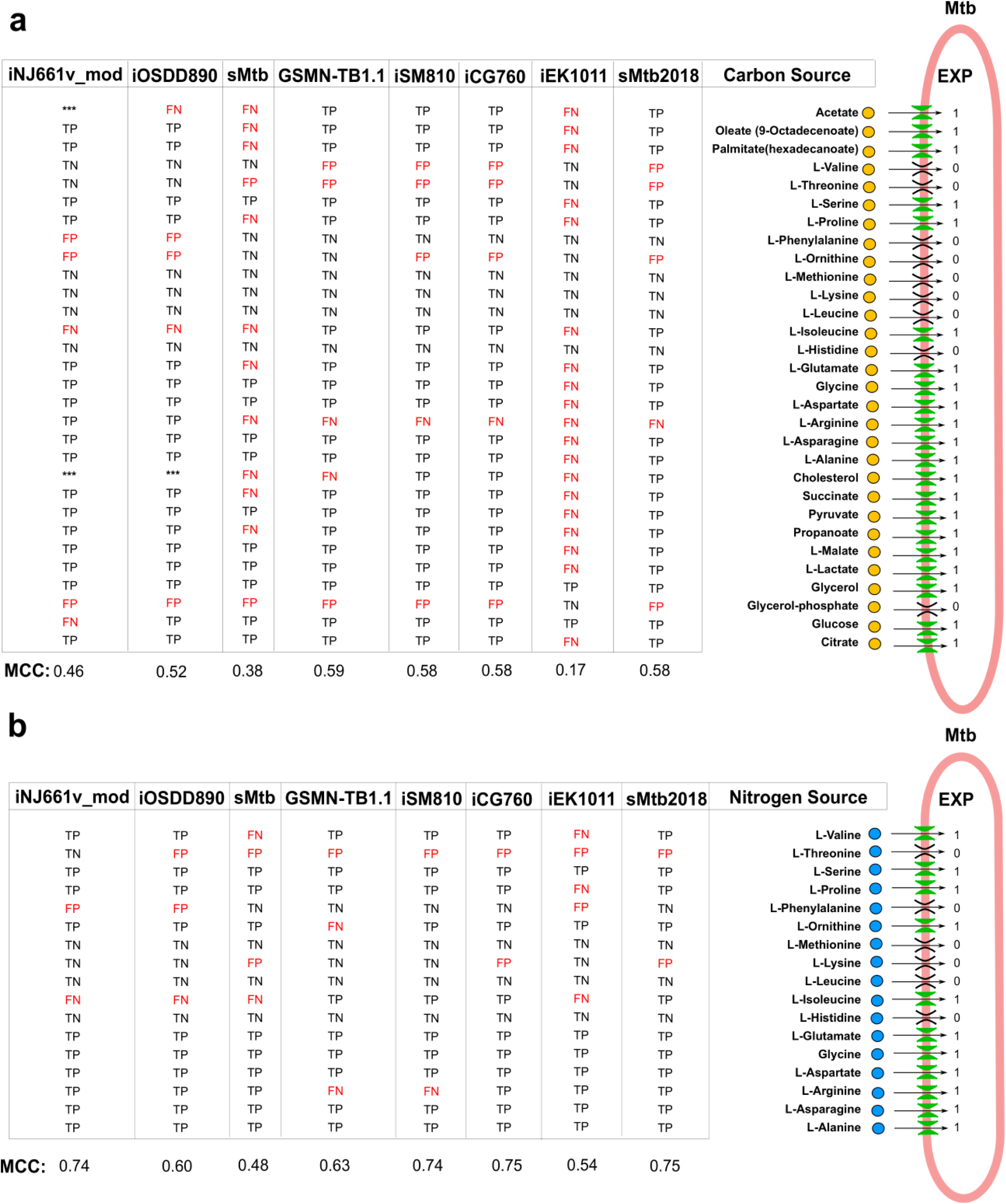
Predictive capacity of Mtb genome-scale models for the utilization of sole carbon and nitrogen sources; **(a)** Growth predictions of Mtb genome-scale models using sole carbon sources; **(b)** Growth predictions of Mtb genome-scale models by using sole nitrogen sources. Model’s performance was evaluated by computation of the Mathews Correlation Coefficient (MCC). Experimental growth data were obtained from [14, 65]. The 1 and 0 values represent growth and not growth in a specific substrate, respectively. Carbon or Nitrogen substrates are classified as TP if model simulation predicts growth while growth is observed experimentally, FP if model simulation predicts growth while no growth is observed experimentally, TN if model simulation predicts no growth while no growth is observed experimentally and FN if model simulation predicts no growth while growth is observed experimentally.

Whilst carbon metabolism has been intensively studied *in vitro* and *ex vivo*, attention has only recently been directed to nitrogen metabolism [72–75]. Similar to carbon consumption, iEK1011 and sMtb were poor at predicting growth on sole nitrogen sources (Fig 4B, Mathews Correlation Coefficient (MCC) = 0.54 and 0.48, respectively). However, like carbon the addition of the menaquinone linked succinate dehydrogenase reaction into iEK1011 and sMtb significantly improves the predictive power of these models on sole nitrogen sources (S16I and S16J Tables). Specifically, the correct growth prediction were obtained for Mtb growing on branched chain amino acids (isoleucine and valine) and proline (Fig 4B). The possible explaination for these correct prediction is that complete degradation of these amino acids can converge on succinate via methyl citrate cycle (degradation of isoleucine and valine) or the GABA shunt (degradation of proline) which couples TCA cycle with oxidative phosphorylation by succinate dehydrogenase.

### Refining Mtb GSMNs

Overall iEK1011 and sMtb2018 were the best overall GSMN’s in terms of genetic background, network topology, number of blocked reactions, mass and charge balance reactions and gene essentiality predictions (Fig 2, Table 2, Table 3, and S1 Fig) and therefore we selected these models to refine further. iEK1011 has the advantage of containing standardized BiGG nomenclature of metabolites and therefore can easily be integrated into the human GSMN Recon3D [76] to simulate intracellular growth, while sMtb2018 has the utility that this model supports *in silico* growth in a wider variety of different nutritional conditions. Our analysis also highlighted some fundemental issues with these models which we addressed in order to improve the performance of these exempler GSMN’s.

As already discussed the new sMtb2018 includes an updated respiratory chain including menaquinone and menaquinol as electron carriers in all respiratory chain reactions and selected ubiquinone-dependent reactions in sMtb were changed to menaquinone-dependent. Six new menaquinone-dependent reactions were included in the sMtb model e.g., succinate dehydrogenase, and cytochrome bc1 menaquinone-dependent, fumarate reductase, and malate dehydrogenase [22]. This improved the predictive growth metric of sMtb and importantly allowed *in silico* growth on cholesterol (see sMtb2018, Fig 4A). Similarly, we added to iEK1011 a menaquinone-dependent succinate dehydrogenase to improve the performance of this model when growing in fatty acids and cholesterol. Further improvements were also made to cholesterol metabolism by updating both models to include the biochemical degradation of the C and D rings of cholesterol which was not known when these models were reconstructed (S17A and S17B Tables) [77].

Although the functionality of the vitamin B12 biosynthetic pathway remains undefined, recent elevated non-synonymous nucleotide substitution rates found on cobalamin-dependent enzymes of 3798 Mtb strains suggest positive selection on these genes and a possible functional biosynthetic pathway [41]. Therefore, we facilitated co-factor metabolism in the models by unblocking B12 synthesis and including the co-factors biotin and pyridoxal-5-phosphate to enhance the phenotype prediction of sMtb2018 and iEK1011 as recommended by Xavier and colleagues [19]. As iEK1011 contains glycerol within its biomass formulation, which prevent *in silico* growth in glycerol-replete media, we incorporated the iNJ661v_mod biomass objective function into this model (S17B Table). We also added additional genes into sMtb2018 based on iOSDD890 genes that were classified as true negatives and true positives by the gene essentiality analysis (S18 Table). Fifty-one reactions that belong to pathways such as glycolysis, gluconeogenesis, TCA cycle, amino acid metabolism and mycolic acid pathway were improved in terms of GPR annotation. All updates for the new models sMtb2.0 and iEK1011_2.0 are summarized in S17 Table.

The sMtb2.0 and iEK1011_2.0 models were curated to remove possible reactions that participate in TICs respectively (methods section). (S19 Table). The stoichiometric matrix associated with these reactions was built and the null space basis vector was computed (S20 Table) [44, 47] identifying twelve TICs within sMtb2.0; (Fig 5A, S2 Appendix) and seven TICs in iEK1011_2.0 (Fig 5B, S3 Appendix); The major TIC of sMtb2.0 were within the reactions required for folate metabolism catalysed by thymidylate synthase (thyA and thyX) enzymes and dihydrofolate reductase (DFRA) enzymes, which is an essential step for *de novo* glycine and purine biosynthesis, and for the conversion of deoxyuridine monophosphate (dUMP) to deoxytimidine monophosphate (dTMP). (Fig 5A). As recommended by Pereira and colleagues [30], we modified the model to use NADPH/NADP^+^ by retaining the DFRA2 and DFRA4 reactions and eliminating the NADH/NAD^+^-dependent reactions, DFRA1 and DFRA3. Our thermodynamic calculations suggested that THYA and THYX (Δ*_r_G_min_* = −123 kJ/mol, Δ*_r_G_max_* =−9.3 kJ/mol and Δ*_r_G_min_* =−160 kJ/mol, Δ*_r_G_max_* =−33 kJ/mol, respectively) are irreversible in the forward direction (S21 Table).

**Figure 5.**
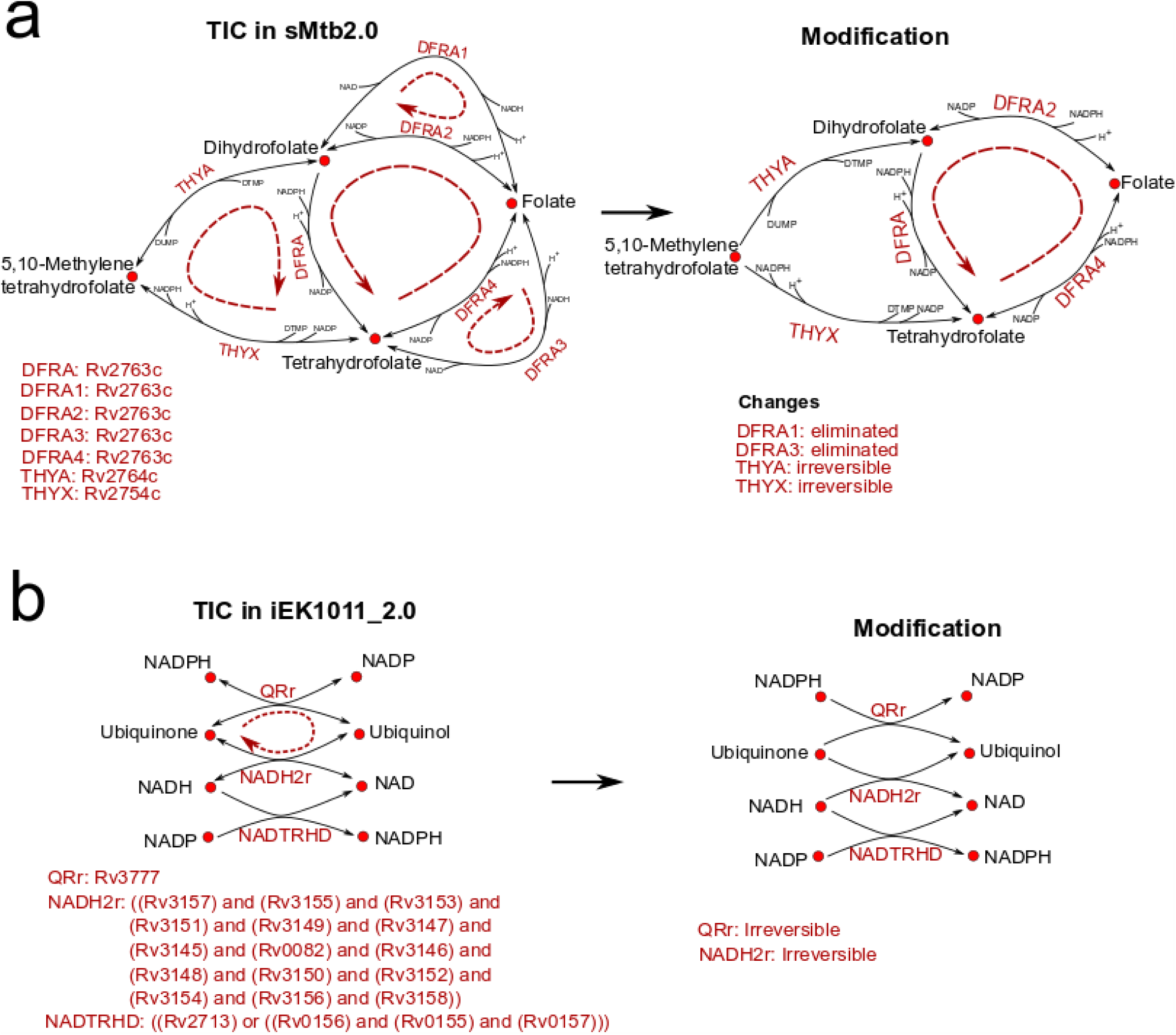
Thermodynamic Infeasible Cycles found in sMtb2.0 and iEK1011_2.0 with proposed modifications; **(a)** TIC at folate metabolism in sMtb2.0; **(b)** TIC affecting ubiquinone oxidoreductases in iEK1011_2.0.

Our analysis showed that two-ubiquinone oxidoreductases (QRr, NADH2r) and a transhydrogenase reaction (NADTRHD) were thermodynamically unstable within the iEK1011_2 as both QRr and NADH2r were reversible. The Gibbs free energy computations suggest these reactions should be unidirectional in the direction of ubiquinol production (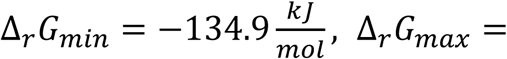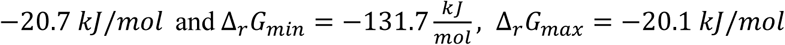, respectively) and therefore we changed the model accordingly to eliminate this TIC.

The sMtb2.0 encompasses 1321 reactions, 1054 metabolites and 989 genes, while iEK1011_2.0 comprises 1237 reactions, 977 metabolites and 1013 genes.

Similar to the above sections, the predictive capability of these modified models was evaluated by simulating gene essentiality predictions using experimental data. In this case, we use the 5% of the specific growth rate as essentiality threshold; if a specific growth rate of no more than 5% of the wild-type is obtained, the given gene was considered as essential, otherwise was considered non-essential (S22 Table). iEK1011_2.0 has the highest predictive performance of gene essentiality in the four media conditions tested (glycerol and cholesterol minimal medium, Middlebrook 7H9, and YM medium) compared with all the Mtb GSMNs evaluated, including sMtb2.0 (8% higher in Cholesterol and Glycerol minimal medium, and 13 % higher in 7H9 medium, Table 4).

**Table 4.** Genome-scale model features for sMtb2.0 and iEK1011_2.0.

| Metrics | sMtb2.0 | iEK1011_2.0 |
| --- | --- | --- |
| **Genes** | 989 | 1013 |
| **Reactions** | 1321 | 1237 |
| **Intracellular Reactions** | 1198 | 1119 |
| **Exchange Reactions** | 122 | 119 |
| **Metabolites** | 1054 | 977 |
| **% of Unbounded Reactions** | (49) 4% | 20 (1.8 %) |
| **MCC Cholesterol minimal medium** | **0.55** | **0.63** |
| **Evaluated Genes** | 834 | 866 |
| True Positive Genes | 209 | 210 |
| True Negative Genes | 449 | 503 |
| False Positive Genes | 58 | 21 |
| False Negative Genes | 118 | 132 |
| **MCC Glycerol minimal medium** | **0.54** | **0.62** |
| **Evaluated Genes** | 834 | 866 |
| True Positive Genes | 207 | 206 |
| True Negative Genes | 449 | 503 |
| False Positive Genes | 58 | 21 |
| False Negative Genes | 120 | 136 |
| **MCC 7H9 medium** | **0.56** | **0.69** |
| **Evaluated Genes** | 984 | 1006 |
| True Positive Genes | 202 | 216 |
| True Negative Genes | 600 | 667 |
| False Positive Genes | 96 | 45 |
| False Negative Genes | 86 | 78 |
| **MCC MtbYM medium** | **0.43** | **0.52** |
| **Evaluated Genes** | 827 | 858 |
| True Positive Genes | 153 | 152 |
| True Negative Genes | 458 | 509 |
| False Positive Genes | 51 | 17 |
| False Negative Genes | 165 | 180 |
| **MCC Unique Carbon Source** | **0.58** | **0.67** |
| **Evaluated Metabolites** | 30 | 30 |
| True Positive Metabolites | 20 | 20 |
| True Negative Metabolites | 5 | 6 |
| False Positive Metabolites | 4 | 3 |
| False Negative Metabolites | 1 | 1 |
| **MCC Unique Nitrogen Source** | **0.75** | **0.75** |
| **Evaluated Metabolites** | 17 | 17 |
| True Positive Metabolites | 11 | 11 |
| True Negative Metabolites | 4 | 4 |
| False Positive Metabolites | 2 | 2 |
| False Negative Metabolites | 0 | 0 |

The ability of these models to predict growth sole carbon and nitrogen sources is also better in the updated GSMN’s as compared with their predecessors. These results show that iEK1011_2.0 and sMtb2.0 are currently the most suitable models for studying host-pathogen interactions, as they are able to correctly predict growth on several important carbon sources that Mtb uses intracellularly [62].

By systematically evaluating eight of the recent Mtb GSMNs, we have identified a number of areas in which model veracity can be improved. Mtb models descended from GSMN-TB (GSMN-TB1.1, iSM810 and iCG760) contain many reactions that are unbalanced by charge and mass, often because protons and water have not been accounted for. We observed that all of the Mtb models have dead-end metabolites particularly in cofactor metabolism and related pathways. The best performing models were identified as sMtb2018 and iEK1011 by virtue of higher gene coverage, fewer blocked reactions, better mass and charge balances, and overall best predictive power of gene essentiality data. Whilst our analysis identified that iEK1011 and sMtb2018 were the overall best performing GSMN’s we also highlighted areas for improvements. For example the iEK1011 model had poor predictive power for growth on sole carbon and nitrogen sources and unlike Mtb cannot grow when cholesterol and fatty acids are provided as sole carbon sources. We thus updated these two models by the addition of new reactions, gap filling of cofactor metabolism, and the identification and curation of TICs, to give sMtb2.0 (derived for sMtb2018) and iEK1011_2.0 (derived from iEK1011).

The improved GSMN’s, sMtb2.0 and iEK1011_2.0 are now available (S1 File) to simulate and predict the metabolic adaptation of Mtb (by the integration of OMICS data) in a plethora of *in vitro* and *in vivo* intracellular conditions. We encourage researchers to continue to curate these models as new data and methods become available. Improved GSMN’s including macrophage-Mtb models provide a platform for more reliable simulations and ultimately a better understanding of the underlying biology of Mtb.

## Methods

All simulations were conducted on a laptop running Windows 10 (Microsoft) using MATLAB 2016a (MathWorks Corporation, Natick, Massachusetts, USA), COBRA Toolbox version 3.0 [33] and Gurobi Optimizer version 7.5.2 (Gurobi Optimization, Inc., Houston, Texas, USA). All code written for this study is available in supplementary information (S2-S5 Files). Genome-scale models of Mtb

Models were obtained from supplementary information of published papers and modified as follows:

- GSMN-TB1.1 – from [14] supplementary info.
- iOSDD890 – from [16] supplementary info.
- sMtb – from [15] supplementary info. Modification included were the addition of exchange reactions to allow constraints by growth medium components.
- iCG760 – from [17] supplementary info.
- iSM810 – from [18] supplementary info.
- iNJ661v_modified – from [19] supplementary info.
- sMtb2018 – from [22] supplementary info.
- iEK1011 – from [24] supplementary info.

### Network connectivity evaluation

GSMNs of Mtb were transformed to substrate networks by local scripts after eliminate biomass reaction. Node-specific topology metrics were carried out using the plugin Network Analyzer [78] in Cytoscape 3.4 [79]. Two main topological parameters were evaluated for each model: 1) the node degree of each metaboliteand 2) the clustering coefficient (S4 Table).

MC3 Consistency Checker algorithm [38] was used to identify Single Connected and Dead End metabolites, and zero-flux reactions in each metabolic network model of Mtb. This algorithm uses a stoichiometric-based identification of metabolites connected only once in each metabolic network and utilize Flux-Variability-Analysis (FVA) for identifying reactions that cannot carry flux [80].

### Biomass Molecular Weight Check

Testing the biomass Molecular Weight consistency was done by running the script of Chan and colleagues [34]. A biomass reaction is not standardized when the Molecular Weight of the biomass formula is not equal to 1 g/mmol. However, the accuracy of the results relies on the correct chemical formulae of metabolites in the tested GSMNs.

### Identification of Unbounded Reactions (URs)

A straightforward way to identify all reactions that participate in one or more TICs is by performing flux variability analysis (FVA). All the Infeasible loops are evidenced as a set of reactions able to carry an unbounded metabolic flux under finite or even zero substrate uptake inputs. The URs are those reactions that by applying FVA [80], their fluxes will hit the values defined by the upper and/or lower bounds constraints [47]. Therefore, we performed FVA with the eight Mtb GSMNs with all the uptake media constraints defined by 1.0 mmol/gDW/h.

### Identification of the core set of TICs

Schellenberger and colleagues [81] used a methodology for identifying the core set of TICs, which form the basis of all such possible cycles. This core can be obtained by the computation of the null space basis of the stoichiometric matrix (all possible thermodynamically infeasible cycles form the null space of the stoichiometric matrix). Consequently, the set containing all the reactions that we previously identified participate in TICs was used to build a stoichiometric matrix. Therefore, the null space basis of this set was computed and the different cycles composed by two and more reactions were identified by a local script.

### Checking the existence of Energy Generating Cycles (EGCs)

Energy generating cycles (EGCs) exist in metabolic networks and can charge energy metabolites like ATP, GTP, CDP, and UTP without any input of nutrients; therefore, their elimination is essential for correcting energy metabolism [49, 82]. Fritzemeier and colleagues developed a methodology for identifying if genome-scale models contain EGCs [49]. We applied it in two steps : 1) Addition of Energy dissipation reactions (EDR) for ATP, GTP, CTP and UTP in the form : H2O[c] + XTP[c] -> H[c] + XDP[c] + Pi[c] and 2) maximization of each EDR flux 𝒗_𝒅_ while no substrate uptake is allowed into the model as follows:

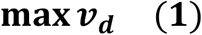

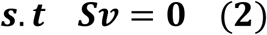

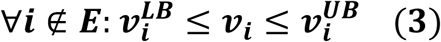

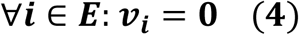

Here, 𝑺 is the stoichiometric matrix, 𝒗 the vector of fluxes, d the index of EDRs, 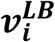 and 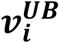 the vector of lower and upper bounds, respectively, and 𝑬 is the set of indices of all exchange reactions of the model. If the optimal value of 𝒗_𝒅_ for this optimization is 𝒗_𝒅_ > 0, there exist in the genome-scale model at least one EGC that is able to generate energy metabolites like ATP, GTP, CTP, or UTP.

### Curation of TICs

Two types of modifications were performed on the curated Mtb genome-scale metabolic network in order to eliminate TICs [47].

i) TICs formed by linearly dependent reversible reactions: Usually, these arise when there are two reactions (NAD^+^- and NADP^+^-dependent) with the same catalytic activity. In this instance, we forced the use of NADPH/NADP^+^ in anabolic reactions and NADH/NAD^+^ for catabolic reactions, as recommended by Pereira and colleague [30]. If two irreversible reactions that catalyze the forward and backward direction exist, both reactions (and GPR rules) are lumped together in just one reversible reaction.
ii) TICs formed by erroneous directionality assignments: we restricted the reaction directionality based on Gibbs free energy change (NExT algorithm) [83, 84] as long as gene essentiality predictions are not compromised

The later modification was based on the utilization of the NExT (network-embedded thermodynamic analysis) algorithm [83, 84]. This algorithm allows identify new irreversible reactions by calculating the thermodynamically feasible range of Gibbs energy of reactions and metabolite concentrations. NExT was implemented for those reactions participating in TICs under Matlab [84] with physiological conditions adapted for Mtb (Table 5). Standard Gibbs energy of formation (Δ***_f_G_i_***) (in kJ/mol), number of hydrogen atoms, and charge of all metabolites involved in TICs were obtained from the Biochemical Thermodynamic Calculator, eQuilibrator [85, 86].

**Table 5.** Biophysical properties and concentration ranges for intracellular Mtb.

| Properties | Values | Reference |
| --- | --- | --- |
| Redox potential, cytosol | -275 mV | Bhaskar et al., 2014 |
| pH, intracellular | 5.7 | Zhang et al., 2003 |
| pH, extracellular (activated macrophage) | 4.5 | Rohde et al., 2007; Vandal et al., 2009 |
| Ionic strength | 0.15 M | Kümmel et al., 2006; Martínez et al., 2014 |
| Oxygen Concentration (mM) | 0.0001 - 0.1 | Martínez et al., 2014 |
| [CO<sub>2</sub>],[Pi] (mM) | 1 - 100 | Kümmel et al., 2006; Haraldsdóttir et al., 2012 |
| Other metabolites (mM) | 0.0001 – 0.1 | Martínez et al., 2014 |
| NADH/NAD | 0.0001 – 0.1 | Martínez et al., 2014 |
| NADPH/NADP | 0.0001 – 0.1 | Martínez et al., 2014 |

If a reaction specified to be reversible in the set of TICs and it has its maximum Δ***_r_***𝑮 calculated to be negative, the reaction is considered to occur in the forward direction. In contrast, if the minimum Δ***_r_***𝑮 is positive, the reaction is considered to occur in the reverse direction. No direction can be inferred when the minimum Δ***_r_***𝑮 is negative and the maximum is positive. Changes in directionality of reactions were done strictly when gene essentiality predictions in the curated genome-scale model were not compromised.

### Gene Essentiality Analysis

To identify essential genes of Mtb grown on individual conditions (cholesterol minimal medium and glycerol minimal medium) from Griffin’s paper [54], we use the Bayesian/Gumbel method of TRANSIT, version 2.02 [59]. The Bayesian/Gumbel method determines posterior probability of the essentiality of each gene (zbar). When zbar value is 1, or close to 1, the gene is considered essential (ES), if zbar is 0, or close to 0, the gene is considered non-essential (NE), uncertain (U) genes are those with zbar values between 0 and 1, and for too small (S) genes zbar is −1. After loading the TA count files (replicates for cholesterol and glycerol) and the gene annotation file into TRANSIT, and running the Gumbel method with default parameters, we obtained an output file with essentiality results (Table S9 and S10). Uncertain (U) and too small (S) genes were not taken into account for the *in silico* essentiality analysis. Minato and colleagues used the same statistical method for classify essential genes [57]. Conversely, DeJesus and colleagues [56] used a Hidden Markov Model based statistical method for classifying genes into four essentiality states: essential (ES), growth defect (GD), nonessential (NE), and growth advantage (GA). In order to evaluate the performance of the Mtb GSMNs to predict gene essentiality data, we use only binary classifiers, therefore we reclassify these genes just in two groups as follows: NE genes included NE and GA genes, and ES genes included GD and ES genes.

For the *in silico* gene essentiality analysis, we set the simulation conditions (asparagine, phosphate, sodium, ammonium, citrate, sulfate, zinc, calcium, chloride, Fe3+, Fe2+, and glycerol or cholesterol) according to Griffin minimal medium [54], 7H9 OADC medium, and “MtbYM” medium and a FBA-based gene essentiality analysis was performed in the eight Mtb models using the “single gene deletion” function of Cobra Toolbox. Default maximization of biomass objective function was used to predict growth in all models. If a specific growth rate of no more than 5% of the wild-type is obtained, the given gene was considered as essential (*in silico*), otherwise was considered non-essential.

Percentage of *in silico* gene essentiality predictions were categorized as: true-positive, false-positive, true-negative, and false-negative when the *in silico* data were compared with experimental essentiality data.

TP (true-positive): model simulation predicts no growth when essential genes are deleted.

FP (false-positive): model simulation predicts no growth when not essential genes are deleted.

TN (true-negative): model simulation predicts growth when not essential genes are deleted.

FN (false-negative): model simulation predicts growth when essential genes are deleted.

For evaluate the performance of the Mtb GSMNs we used sensitivity, specificity, accuracy, and Mathews correlation coefficient (MCC) metrics:

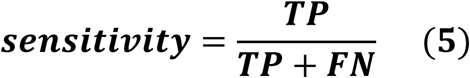

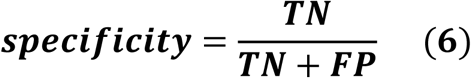

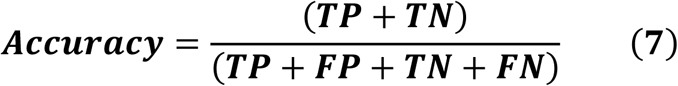

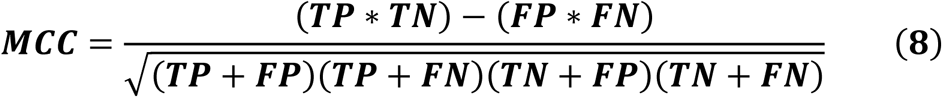

### Utilization of carbon and nitrogen sources

The methodology for modeling the effect of different carbon sources and nitrogen sources on Mtb growth was adapted from Lofthouse and colleagues [14]. The biology Phenotype MicroArray experiments classification used were obtained from Lofthouse and colleagues, 2013. They classified growth and no-growth in different carbon and nitrogen sources from the original Biolog data of Khatri and colleagues and the Roisin’s minimal media [14, 65]. In addition, we used Roisin’s minimal media data that also were obtained by Lofthouse and colleagues.

To model the carbon source experiment, we simulated the media as a modified form of Roisin’s minimal media containing unlimited quantities of ammonia, phosphate, iron, sulfate, carbon dioxide and a Biolog carbon source influx of 1 mmol/gDW/h. Similarly, the nitrogen source experiment was simulated using a modified form of Roisin’s media, where ammonia was replaced with 1 mmol/g DW/h of the Biolog nitrogen source and pyruvate was used as a carbon source (influx at 1 mmol/g DWt/h).

For comparing the utilization of carbon and nitrogen sources in all Mtb models with experimental data, we used Mathew’s correlation coefficient metrics (equation 8).

*In silico* growth predictions in carbon and nitrogen sources also were categorized as: true-positive, false-positive, true-negative, and false-negative.

TP (true-positive): model simulation predicts growth while growth is observed experimentally (or respiration rate is observed in biolog phenotype microarrays) in presence of a unique carbon or nitrogen source.

FP (false-positive): model simulation predicts growth while no growth is observed experimentally in presence of a unique carbon or nitrogen source.

TN (true-negative): model simulation predicts no growth while no growth is observed experimentally in presence of a unique carbon or nitrogen source.

FN (false-negative): model simulation predicts no growth while growth is observed experimentally in presence of a unique carbon or nitrogen source.

## Supporting Information

**S1 Appendix. Overview of the eight constraint-based models of Mtb used in this study.** A document providing additional detail about the eight Mtb GSMNs used in this study. (DOCX)

**S2 Appendix. Curation of additional Thermodynamic Infeasible Cycles in sMtb2.0.** A document providing detailed description of the curation of TICs in sMtb2.0. (DOCX)

**S3 Appendix. Curation of additional Thermodynamic Infeasible Cycles in iEK1011_2.0.** A document providing detailed description of the curation of TICs in iEK1011_2.0. (DOCX)

**S1 Fig. ROC curves of gene essentiality predictions for Mtb GSMNs**. **a** Receiver operating characteristic curve for the gene essentiality predictions in cholesterol minimal medium, **b** Receiver operating characteristic curve for the gene essentiality predictions in glycerol minimal medium, **c** Receiver operating characteristic curve for gene essentiality predictions in 7H9 Middlebrook OADC medium, **d** Receiver operating characteristic curve for gene essentiality predictions in MtbYM medium. (TIF)

**S2 Fig. central role of succinate dehydrogenase in oxidation of odd-chain substrates. (TIF)**

**S1 Table. Set analysis of genes from all the eight Mtb GSMNs.** A table with pairwise comparisons, unions, and intersections of genes annotated in the eight GSMNs of Mtb. (XLSX)

**S2 Table. Metabolic pathways associated to all the intersected gene sets of Mtb GSMNs. (XLSX)**

**S3 Table. Genes and metabolic pathways shared between GSMNs of Mtb.** (XLSX)

**S4 Table. Network topological properties of Mtb GSMNs. (**XLSX)

**S5 Table. List of unbalanced reactions and Metabolites without formulas in the Mtb GSMNs.** (XLSX)

**S6 Table. List of blocked reactions and dead-end metabolites. (XLSX)**

**S7 Table. List of Thermodynamically infeasible cycles identified for the eight Mtb GSMNs.** (XLSX).

**S8 Table**. **Transposon sequencing analysis of Mtb genes required for growing on minimal medium plus cholesterol.** A list of genes of Mtb classified as essential, non-essential, and uncertain for growing in cholesterol minimal medium, the essentiality analysis was obtained by applying the Bayesian/Gumbel Method incorporated into the software TRANSIT. (XLSX)

**S9 Table. Transposon sequencing analysis of Mtb genes required for growing on minimal medium plus glycerol.** A list of genes of Mtb classified as essential, non-essential, and uncertain for growing in glycerol minimal medium, the essentiality analysis was obtained by applying the Bayesian/Gumbel Method incorporated into the software TRANSIT. (XLSX)

**S10 Table. Predictive power of Mtb GSMNs for classify essential and non-essential genes on cholesterol minimal medium.** Genes whose *in silico* knockouts give growth rate values lesser than 5% of the maximum growth rate are classified as essential, otherwise are classified as non-essential. (XLSX)

**S11 Table. Predictive power of Mtb GSMNs for classify essential and non-essential genes on glycerol minimal medium.** Genes whose *in silico* knockouts give growth rate values lesser than 5% of the maximum growth rate are classified as essential, otherwise are classified as non-essential. (XLSX)

**S12 Table. Predictive power of Mtb GSMNs for classify essential and non essential genes on 7H9 OADC medium.** Genes whose *in silico* knockouts give growth rate values lesser than 5% of the maximum growth rate are classified as essential, otherwise are classified as non-essential. (XLSX)

**S13 Table. Predictive power of Mtb GSMNs for classify essential and non-essential genes on cholesterol minimal medium.** Genes whose *in silico* knockouts give growth rate values lesser than 5% of the maximum growth rate are classified as essential, otherwise are classified as non-essential. (XLSX)

**S14 Table. List of common False Positive and False Negative Genes of all the Mtb GSMNs during gene essentiality predictions**. (XLSX)

**S15 Table. Predictive power of Mtb GSMNs growing on sole carbon sources.** (XLSX)

**S16 Table. Predictive power of Mtb GSMNs growing on sole nitrogen sources.** (XLSX)

**S17 Table. New added reactions into the sMtb and iEK1011 models.** (XLSX)

**S18 Table. List of reactions with modified gene annotation in sMtb2.0.** (XLSX)

**S19 Table. List of reactions that participate in Thermodynamically Infeasible Cycles.** (XLSX)

**S20 Table. Null Space of the stoichiometric matrix formed by unbounded reactions of sMtb2.0 and iEK1011_2.0.** (XLSX)

**S21 Table. Gibbs free energy change of Unbounded Reactions of updated Mtb GSMNs.** (XLSX)

**S22 Table. Gene Essentiality Predictions for iEK1011_2.0 and sMtb2.0 on four Mtb media.** (XLSX)

**S23 Table. Growth Phenotypes of iEK1011_2.0 and sMtb2.0 on unique carbon and nitrogen sources.** (XLSX)

**S1 File. Updated Mtb models sMtb2.0 and iEK1011 in .mat and, xlsx format.** (ZIP)

**S2 File. Matlab script for checking the charge and mass balance of Mtb GSMNs.** (ZIP)

**S3 File. Matlab scripts for identifying unbounded reactions and thermodynamically infeasible cycles in Mtb GSMNs.** (ZIP)

**S4 File. Matlab scripts for identifying Mtb GSMNs with Energy Generating Cycles.** (ZIP)

**S5 File. Matlab scripts for gene essentiality analysis of Mtb GSMNs on four media conditions.** (ZIP)

## Author Contributions

**Conceptualization:** Víctor A. López-Agudelo, Dany JV Beste., Rigoberto Rios-Estepa.

**Data curation:** Víctor A. López-Agudelo

**Formal analysis:** Víctor A. López-Agudelo, Emma Laing, Tom Mendum, Dany JV. Beste, Rigoberto Rios-Estepa.

**Funding acquisition:** Dany JV. Beste, Rigoberto Rios-Estepa.

**Investigation:** Víctor A. López-Agudelo, Emma Laing, Tom Mendum, Andres Baena, Luis F. Barrera, Dany JV. Beste, Rigoberto Rios-Estepa.

**Methodology:** Víctor A. López-Agudelo, Dany JV. Beste, Rigoberto Rios-Estepa.

**Project administration:** Rigoberto Rios-Estepa, Dany JV. Beste.

**Resources:** Víctor A. López-Agudelo, Emma Laing, Tom Mendum, Andres Baena, Luis F. Barrera, Dany JV. Beste, Rigoberto Rios-Estepa.

**Software:** Víctor A. López-Agudelo, Tom Mendum, Dany JV. Beste.

**Supervision**: Dany JV. Beste, Rigoberto Rios-Estepa.

**Validation**: Tom Mendum, Dany JV. Beste, Rigoberto Rios-Estepa.

**Visualization:** Víctor A. López-Agudelo

**Writting – original draft:** Víctor A. López-Agudelo, Andres Baena, Luis F. Barrera, Dany JV. Beste, Rigoberto Rios-Estepa

**Writting – review & editing:** Víctor A. López-Agudelo, Tom Mendum, Dany JV. Beste, Rigoberto Rios-Estepa.

## Acknowledgments

Víctor A. López-Agudelo acknowledges the economic support provided by the Administrative Department of Science, Technology and Innovation - COLCIENCIAS, Colombia, during his Ph.D. studies (National Ph.D. scholarship Conv. 727-2015).

